# Unbiased association and expression studies identify novel genes for tooth development

**DOI:** 10.1101/209684

**Authors:** Meredith A. Williams, Claudia Biguetti, Miguel Romero-Bustillos, Kanwal Maheshwari, Nuriye Dinckan, Franco Cavalla, Xiaoming Liu, Renato Silva, Sercan Akyalcin, Z Oya Uyguner, Alexandre R. Vieira, Brad A. Amendt, Walid D Fakhrouri, Ariadne Letra

## Abstract

Previously reported co-occurrence of colorectal cancer (CRC) and tooth agenesis (TA) and the overlap in disease-associated gene variants suggest involvement of similar molecular pathways. In this study, we took an unbiased approach and tested genome-wide significant CRC-associated variants for association with isolated TA. Thirty single nucleotide variants (SNVs) in CRC-predisposing genes/loci were genotyped in a discovery dataset composed of 440 individuals with and without isolated TA. Genome-wide significant associations were found between TA and *DUSP10* rs6687758 (P=1.25 × 10^−9^) and *ATF1* rs11169552 (P=4.36 × 10^−10^), with strong association found with *CASC8* rs10505477 (P=8.2 × 10^−5^). Additional CRC marker haplotypes were also significantly associated with TA (P<0.0002). Genotyping an independent dataset consisting of 52 cases with TA and 427 controls confirmed the association with *CASC8*.

Atf1 and Dusp10 expression was detected in the mouse developing teeth from early bud stages to the formation of the complete tooth, suggesting a potential role for these genes and their encoded proteins in tooth development. Our findings suggest Atf1 and Dusp10 as new tooth development genes, while having a role in colorectal cancer. While their individual contributions in tooth development remain to be elucidated, these genes may be considered additional candidates to be tested in future human genetic studies.

## Introduction

Colorectal cancer (CRC) is the third most commonly diagnosed cancer in the world and a leading cause of cancer-related deaths.(Navarro et al. 2017) Its etiology is multifactorial and ~33% of all cases are attributed to genetic factors.(Siegel et al. 2014) Tooth agenesis (TA), the congenital absence of one or more permanent teeth, results from disturbances during the initiation stage of tooth development, and represents one of the most common craniofacial anomalies in humans.(Hennekam 2010) Over a decade ago, Lammi et al.(Lammi et al. 2004) reported that mutations in the tumor suppressor gene *AXIN2* were found co-segregating with colorectal cancer and tooth agenesis in a large multiplex family. Moreover, these authors showed the expression of *AXIN2* in developing mouse teeth and suggested that genes involved in colorectal cancer could also be involved in tooth development. (Lammi et al. 2004)

Tooth development requires a sophisticated series of signaling interactions between the oral epithelium and mesenchyme under strict genetic control by a number of signaling molecules and their downstream signaling pathways.(Thesleff and Sharpe 1997) Any alteration of the epithelial-mesenchymal interactions can have deleterious effects on tooth development, and may affect growth, differentiation and pattern formation, or even have systemic effects.(Cudney and Vieira 2012; Yin and Bian 2015) TA can occur as part of a syndrome, although it is more frequently found as an isolated trait that may appear sporadically or segregating in families, for which the prevalence ranges between ~2-10% excluding third molars.(Polder et al. 2004) Based on the number of missing teeth, TA is referred to as hypodontia (up to 5 teeth missing), oligodontia (≥6 teeth missing), or anodontia (all teeth missing), the latter being mostly associated with syndromic TA.(Hennekam 2010)

Studies in mice have revealed more than 200 genes involved in tooth development and shown the functional importance of *Msx1, Pax9, Pitx2,* and *Lef1* genes in proper tooth formation and morphogenesis, since the absence of these genes results in arrest of tooth development at the bud stage.(Thesleff et al. 1996) In humans, the best characterized mutations involve *MSX1, PITX2,* and *PAX9.*(Li et al. 2014; Thesleff et al. 1996) Additional mutations in Wnt pathway genes (i.e., *AXIN2, LRP6, WNT10A, WNT10B*) have been increasingly implicated in the susceptibility to TA.(Yin and Bian 2015) Intriguingly, it is well established that Wnt pathway genes also play critical roles in tumorigenesis, particularly in colorectal cancer. (Callahan et al. 2009; Lammi et al. 2004; Letra et al. 2009; Yin and Bian 2016)

In addition to epithelial-mesenchymal interactions, cell growth and cell differentiation, the signaling pathways involved in tooth development (i.e., WNT, BMP, SHH, FGF, TGF-β, and NF-kB), also overlap with those implicated in colorectal cancer.(Cudney and Vieira 2012; Yin and Bian 2015) Recently, genome-wide association studies (GWAS) have uncovered over 50 loci and single nucleotide polymorphisms (SNPs) that are significantly correlated with the susceptibility to CRC, most of which map to genes with established roles in tumorigenesis, or involved in developmental processes such as transcriptional regulation, genome maintenance, and cell growth and differentiation.(Hsu et al. 2015; Kang et al. 2015; Zhang et al. 2014) Of note, SNPs in *CDH1, BMP2, BMP4,* and *GREM1* genes, known to have important roles in craniofacial and/or tooth development, (Cudney and Vieira 2012) were also reported as highly associated with CRC.(Cudney and Vieira 2012; Zhang et al. 2014)

Given the previously reported co-occurrence of CRC and TA and the overlap in disease-associated pathways, we sought to identify new genes for TA by testing the association of genome-wide associated CRC variants with isolated TA. Here, we describe the identification of new candidate genes for TA by the combination of an unbiased association study as well as expression analyses in oral and dental tissues of mouse embryos.

## Materials and Methods

### Study Population

This study was approved by the University of Texas Health Science Center at Houston Committee for the Protection of Human Subjects and the University of Pittsburgh Institutional Review Board. Unrelated individuals with and without tooth agenesis were invited to participate in the study when presenting for treatment at the University of Texas Health Science Center at Houston School of Dentistry clinics. Written informed consent was obtained from all participating individuals and/or from their parents or guardians. Clinical and demographic information, and saliva samples were collected from all individuals. When available, individual and familial history of cancer was also recorded. The presence of tooth agenesis was determined by a dentist through clinical and radiographic examinations, and considered if one or more permanent teeth were missing from the oral cavity excluding third molars. Individuals showing signs of syndromic forms of tooth agenesis (i.e., ectodermal dysplasia features) were excluded. To avoid potential effects of population stratification, only individuals with self-reported Caucasian ethnicity were included in the study. A total of 440 unrelated individuals, 93 with nonsyndromic TA (29 males, 64 females) and 347 control individuals without TA or family history of TA (101 males, 246 females) were included in this study. Of the 93 cases with TA, 61 presented with hypodontia while 32 presented with oligodontia. Positive family history of cancer was reported by 28 cases and 33 control individuals (Supplementary Material, Table S1).

### Selection of CRC-Associated Variants and Genotyping

We selected 30 CRC-risk-associated SNPs that reached genome-wide significance (5 × 10-8) in previously published GWAS (Houlston et al. 2010; Hsu et al. 2015; Lemire et al. 2015) to genotype in this study (Table 1). For regions where multiple SNPs have been reported, we included the most significantly-associated SNP according to the Genetics and Epidemiology of Colorectal Cancer Consortium (GECCO; https://share.fhcrc.org/sites/gecco/), and prioritized additional SNPs for genotyping based on: 1) location in regulatory or enhancer regions of a given gene, 2) having an effect on motifs in the distal regulatory regions of respective genes, and/or 3) interactions with known genes of biological relevance.

**Table 1.**
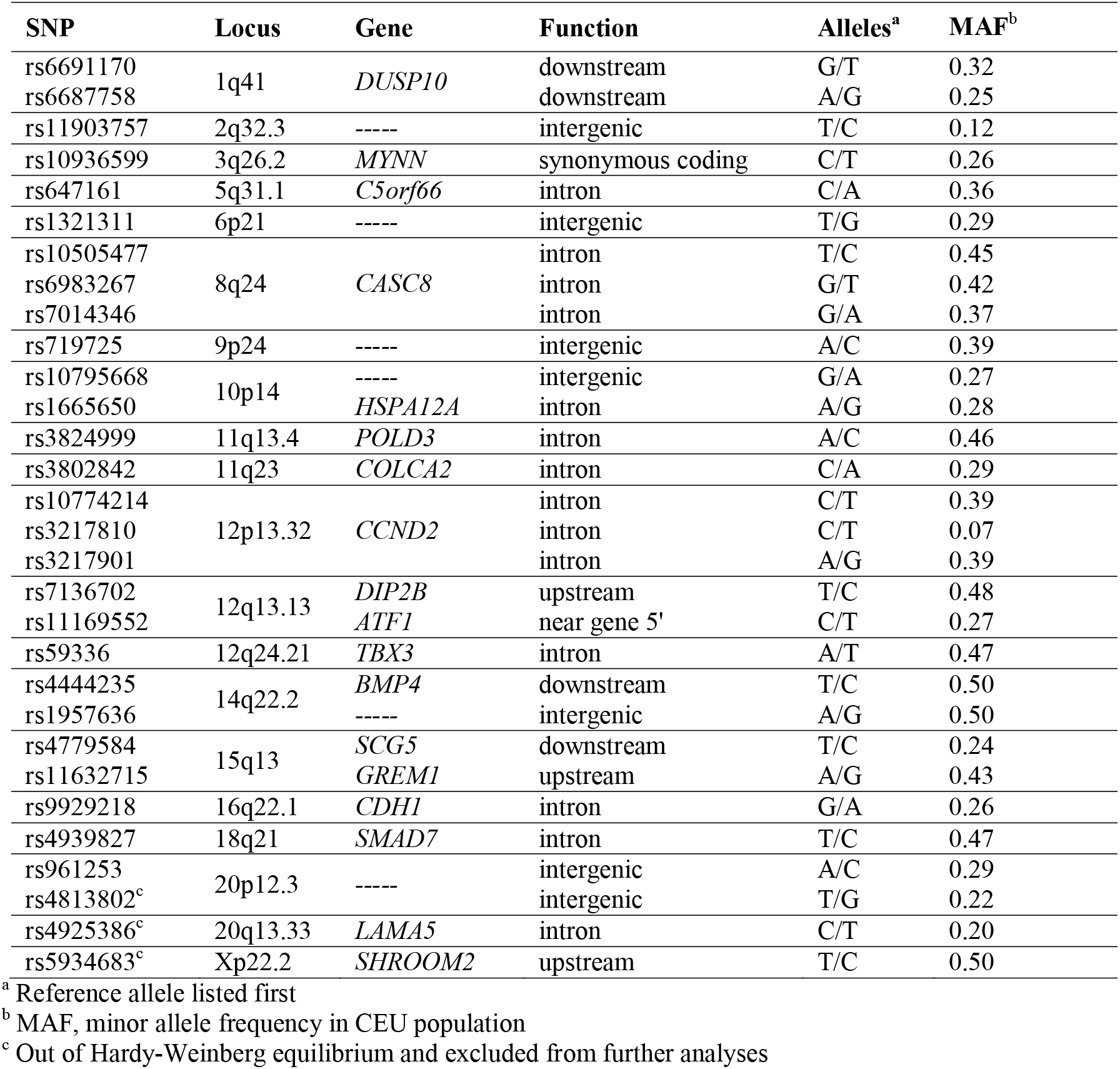
Details of SNPs investigated in this study.

Genotyping was performed using Taqman chemistry (Ranade et al. 2001) in 5-μL final reaction volumes in a ViiA7 Sequence Detection System (Applied Biosystems, Foster City, CA). Results were analyzed using EDS v.1.2.3 software (Applied Biosystems). For quality control of genotyping reactions, we used a non-template reaction as negative control and a DNA sample of known genotype as positive control. The call rate of genotyped SNPs was considered as >95% accurate.

### Data analyses

Power calculations were performed using Genetic Power Calculator (http://zzz.bwh.harvard.edu/gpc/) and indicate that the study sample size provided approximately 86% power to detect an association with an alpha of 0.05, if the markers selected are in linkage disequilibrium with the causal factor (D’=0.80) and their frequencies are around 80%. Data analysis was performed using PLINK version 1.06.(Purcell et al. 2007) Hardy-Weinberg equilibrium was calculated for cases and controls, and SNPs showing evidence of deviation in controls were excluded from further analyses. Differences in allele and genotype frequencies for each polymorphism between cases and controls were compared using chi-square and Fisher Exact tests. We corrected for multiple testing using the Bonferroni method, and the significance level was set considering the number of tests (n=30) to give a corrected P-value (α=0.002). Haplotype analyses were performed using the ‘haplotype-based case-control association test’ as implemented in PLINK. Regression analyses were performed using EpiInfo v.7.1 (https://www.cdc.gov/epiinfo/) to identify potential preferential associations between specific SNP genotypes with tooth agenesis adjusted for family history of cancer and gender.

### In silico prediction of SNP function

Annotation of the newly identified loci was performed using the dbNSFP (Liu et al. 2016) and FuncPred (https://snpinfo.niehs.nih.gov/snpinfo/snpfunc.html) databases. Information on the function of the associated SNPs regarding splicing regulation, stop codon, Polyphen predictions, transcription factor binding site predictions, miRNA binding site prediction regulatory potential score, conservation score, and nearby genes, were recorded.

### Immunofluorescence and immunohistochemistry

The expression of ATF1 and DUSP10 proteins was evaluated using immunofluorescence and immunohistochemistry in developing mouse tooth sections, respectively. Wild type C57BL/6 mouse embryos at embryonic days (E) 12.5, 14.5, 16.5, 18.5 and postnatal day 0 (P0) were harvested, fixed in 4% paraformaldehyde and embedded in paraffin for sectioning at 7μm of thickness. Sections were deparaffinized in xylene solution, rehydrated in gradative alcohol baths and rinsed with deionized H2O at room temperature. For antigen retrieval, sections were immersed in sodium citrate buffer (10mM, pH 6) at 100°C for 30 minutes and allowed to cool down for 30 min at room temperature. To avoid nonspecific binding of the antibodies, sections were immersed in blocking solution (1% bovine serum albumin and 10% goat serum diluted inPBS) and incubated with AffiniPure Fab fragment goat anti-mouse (Jackson ImmunoResearch Laboratories, PA, USA), for 1 hour at room temperature. For DUSP10, an additional blocking step consisted of incubating sections in 2.5% normal horse serum (Vector Laboratories, CA, USA) for 30 min at room temperature. Sections were then washed and incubated with either Atf1 (rabbit polyclonal anti-mouse Atf1; ab189311, Abcam, MA, USA), or Dusp10 (rabbit polyclonal anti-mouse Dusp10; ab71309, Abcam, MA, USA) primary antibodies at 4^o^ C overnight. For Atf1, sections were then washed and incubated with Alexa Fluor^®^ 555 (goat anti-rabbit secondary antibody, A-21428, ThermoFisher Scientific, MA, USA) for 2 hours in a dark chamber at room temperature. Lastly, sections were washed (3 × 10’), counterstained with DAPI (Invitrogen) in a dark chamber for 10 min, and mounted with ProLong Gold AntifadeReagent (Invitrogen). For Dusp10, sections were incubated with biotinylated horse anti-rabbit IgG (BP-1100, Vector Laboratories, CA, USA) for 30 minutes at room temperature, washed and incubated with Streptavidin/Peroxidase complex (PK7800, Vector Laboratories, CA, USA) for 5 minutes and washed. Sections were then incubated in 3,3'-Diaminobenzidine (ImPACT DAB, Vector Laboratories, CA, USA) chromogen for 2 minutes, washed, and counterstained with Mayer’s Hematoxylin. Lastly, sections were washed, dehydrated and mounted.Imaging was performed after 48 hours in a Nikon Eclipse Ni-U upright fluorescence microscope (Nikon, Germany) equipped with a Zyla 5.5 sCMOS camera (Andor).

### Reverse transcriptase-PCR (RT-PCR)

Total RNA was extracted from SCAP by using the RNeasy kit (Qiagen Inc, Valencia, California, USA). RNA sample integrity was checked by analyzing 1 μg of total RNA on 2100 Bioanalyzer (Agilent Technologies, Santa Clara, California, USA). After RNA extraction, complementary DNA was synthesized by using 3 μg of RNA through a reverse transcription reaction using QuantiTectRT kit (Qiagen Inc, Valencia, California, USA). Four sets of primers for *CASC8* amplification were designed using Primer3 (http://bioinfo.ut.ee/primer3-0.4.0/primer3/) (Supplementary Material, Table S4). β-actin (Actb) was used as endogenous control. PCR reactions were performed in 25uL final reaction volume using 20ng of SCAP cDNA, 1uL of each forward and reverse primers, 0.5uL Taq DNA polymerase, MgCl2, deoxynucleotide mix, 10X PCR buffer, and water. Reaction conditions were: 40 cycles at 95°C (10’), 94°C (1’), 56°C (1’), and 72°C (2’). Products were resolved in agarose gel electrophoresis and imaged using Odyssey Scanner (LI-COR Biosciences, Lincoln, NE).

## Results

### GWAS-based association study

A total of 440 unrelated individuals, 93 with nonsyndromic TA (29 males, 64 females) and 347 control individuals without TA or family history of TA (101 males, 246 females) were included in this study. The presence of tooth agenesis was determined through clinical and radiographic examinations, and considered if one or more permanent teeth were missing from the oral cavity excluding third molars. Individuals showing signs of syndromic forms of tooth agenesis (i.e., ectodermal dysplasia features) were not included in the study. Of the 93 cases with TA, 61 presented with hypodontia while 32 presented with oligodontia. Positive family history of cancer was reported by 28 cases and 33 control individuals.

We selected 30 CRC-predisposing SNVs with genome-wide significance (5 × 10^−8^) from previously published GWAS (Houlston et al. 2010; Hsu et al. 2015; Lemire et al. 2015), for genotyping in our case-control dataset (Table 1). For regions where multiple SNPs have been reported, we included the most significantly-associated SNP according to the Genetics and Epidemiology of Colorectal Cancer Consortium (GECCO; https://share.fhcrc.org/sites/gecco/), and prioritized additional SNPs for genotyping based on: 1) location in regulatory or enhancer regions of a given gene, 2) having an effect on motifs in the distal regulatory regions of respective genes, and/or 3) interactions with known genes of biological relevance. Genotypes were generated using Taqman chemistry (Ranade et al. 2001) and automatic call rate was >98%.

Genome-wide significant associations were found between TA and individual SNPs on 1q41 and 12q13 (Table 2). At the 1q41 locus, rs6687758 is located downstream of the dual specificity phosphatase 10 (*DUSP10)* gene, and allelic (P= 2 × 10^−9^) and genotypic (P= 1.25 × 10^−9^) associations were found between this SNP and TA, particularly under a recessive model (P= 1.27 × 10^−9^). At the 12q13.1 locus, rs11169552 is located 2Kb upstream of the activating transcription factor 1 (*ATF1)* gene promoter, for which genome-wide genotypic (P= 4.36 × 10^−10^) and significant allelic associations (P=1.87 × 10^-^7) were also found (Table 2). Strong association was also found between intronic variants in the cancer susceptibility 8 (*CASC8)* gene on chromosome 8q24 and TA for SNP rs10505477 genotypes and alleles (P= 8.16 × 10^−5^ and P= 1.7 × 10^−5^, respectively) and rs7014346 (P=0.0005 for genotype under a recessive model) (Table 2). When stratifying analyses by cases with hypodontia or oligodontia phenotypes, these associations remained significant for both phenotypes, despite the smaller number of oligodontia cases (P ≤ 0.002, data not shown). Additional significant haplotype associations were also found between markers in the associated genes with the risk of isolated TA (Supplementary Material, Table S2). The strongest association was observed for the G-A-G allele haplotypes from *CASC8* SNPs rs10505477, rs7013436, and rs6983267 (P= 5 × 10^−13^), followed by the G-G haplotype from *DUSP10* rs6687758 and rs6691170 (P = 7.32 × 10^−10^). Additional nominal associations were found for *ATF1* haplotypes (P≤ 0.009) (Table S2). Interestingly, significant associations were found between the ‘TT’ genotype for *ATF1* rs11169552 (P=0.00004) and the ‘GG’ genotype for *DUSP10* rs6687758 (P=0.003) with TA phenotypes in the presence of positive family history of cancer (Supplementary Material, Table S3).To confirm our findings, we genotyped the associated SNPs on an independent dataset composed of 52 cases with tooth agenesis and 427 gender- and ethnicity-matched unrelated controls, and found significant association between *CASC8* rs10505477 (P=0.006 for genotype and P=0.008 for allele) and TA, particularly under a dominant model (P= 0.001).

**Table 2.** Summary of association results under different genetic test models.

| Gene | SNP Id. | Test | Alleles<sup>a</sup> | Frequency Cases | Frequency Controls | P-value<sup>b</sup> |
| --- | --- | --- | --- | --- | --- | --- |
| *ATF1* | rs11169552 | Genotype | CC/CT/TT | 38/31/24 | 195/129/14 | **4.36 x 10<sup>-10</sup>** |
|  |  | Allele | CC/TT | 107/79 | 519/157 | **1.85 x 10<sup>-7</sup>** |
|  |  | Dominant | CC x CT + TT | 38/55 | 195/143 | 0.004 |
|  |  | Recessive | CC + CT x TT | 69/24 | 324/14 | **6.79 x 10<sup>-11</sup>** |
| *DUSP10* | rs6687758 | Genotype | AA/AG/GG | 37/34/22 | 217/112/14 | **1.25 x 10<sup>-9</sup>** |
|  |  | Allele | AA/GG | 108/78 | 546/140 | **2.09 x 10<sup>-9</sup>** |
|  |  | Dominant | AA x AG + GG | 37/56 | 217/126 | **5.09 x 10<sup>-5</sup>** |
|  |  | Recessive | AA + AG x GG | 71/22 | 329/14 | **1.27 x 10<sup>-9</sup>** |
| *CASC8* | rs10505477 | Genotype | TT/TC/CC | 16/45/32 | 119/167/56 | **8.16 x 10<sup>-5</sup>** |
|  |  | Allele | TT/CC | 77/109 | 405/279 | **1.69 x 10<sup>-5</sup>** |
|  |  | Dominant | TT x TC + CC | 16/77 | 119/223 | 0.001 |
|  |  | Recessive | TT + TC x CC | 61/32 | 286/56 | 0.0001 |
|  | rs7014346 | Genotype | GG/GA/AA | 31/38/24 | 145/154/39 | 0.002 |
|  |  | Allele | GG/AA | 100/86 | 444/232 | 0.003 |
|  |  | Dominant | GG x GA + AA | 31/62 | 145/193 | 0.09 |
|  |  | Recessive | GG + GA x AA | 69/24 | 299/39 | 0.0005 |

<sup>a</sup> Reference allele listed first
<sup>b</sup> Fisher Exact test, denotes genome-wide significance if $P \leq 5 \times 10^{-8}$ (bolded), and significant under Bonferroni correction if $P \leq 0.002$ (italicized).

### Functional annotation of associated variants

As a first step into assessing the relevance of the associated genes in tooth agenesis phenotypes, we annotated the associated variants using the dbSNP database (Liu et al. 2016) to predict gene and variant function. Our analyses showed that the associated variants, although common, are predicted to have effects on chromatin structure and RNA polymerase activity and on transcription of their respective genes. *ATF1* rs11169552 is located in the gene promoter in a region showing high DNase I hypersensitivity clusters and potential location of regulatory elements (enhancers, silencers, insulators, and locus control region) (Fig. S4). Further, *ATF1* rs11169552 is predicted to harbor a binding site for *POLR2A,* essential for RNA polymerase activity and DNA transcription. *DUSP10* rs6687758 is located in a DNAse I hypersensitivity cluster with binding motif for TBP (TATA-box binding protein) (Fig. S5). *CASC8* rs10505477 appears to harbor a putative binding motif to NK2-5, in a region of enriched H3K27Ac histone marks (Supplementary Material, Fig. S6). These findings suggest that these variants are likely to have functional consequences on gene expression.

### Expression analyses

Since no information was available from public databases regarding the expression of *ATF1, DUSP10 and CASC8* during tooth development, as a second step into determining the relevance of these genes in tooth development, we sought to assess whether their transcripts and/or encoded proteins were expressed in the oral and dental tissues of developing mouse embryos.

We performed immunofluorescence and immunohistochemistry procedures to detect the localization of Atf1 and Dusp10, respectively, during murine tooth development stages. At E12.5, Atf1 expression was detected in the oral epithelial cells and in the subjacent condensed ectomesenchymal cells (Figures 1A and 1C). Later, at E14.5, Atf1 expression was evident in the oral and tongue epithelia, and underlying mesenchyme, in the inner dental epithelium (IE) and in few scattered cells of the dental papilla (DP) (Fig. 1, D-E). At E16.5 (Supplementary Material, Fig. S2), and E18.5, Atf1 was markedly expressed in the inner epithelium and the stratum intermedium. Reduced expression was also noted in the cells from the stratum intermedium, stellate reticulum and outer dental epithelium (Figures 1F and 1H). At P0, Atf1 expression shifted to the cytoplasm of the polarized layer of ameloblasts, in a perinuclear pattern in the ameloblast bodies, as well as in the Tomes’ processes (Figures 1I and 1J). *Atf1* mRNA expression had been shown to be expressed in mouse embryos as early as E10.5, with strong expression noted in the mesenchyme of frontonasal prominences, branchial arches and limbs (Gray et al. 2004) (Figure 1K). Dusp10 expression was detected at a site of proliferation of the dental placode at E12.5, surrounded by a condensation of the underlying ectomesenchymal cells compatible with a bud stage of tooth development. Expression was evident in the epithelial cells from the oral epithelium, as well as in the condensed ectomesenchymal cells surrounding the epithelial proliferation (Figures 2A and 2B). At E14.5, nuclear and cytoplasmic staining was observed on the cells from the enamel organ and dental papillae (Figures 2C and 2D). At bell stage E16.5, positive staining was observed in the enamel organ, dental papillae and dental follicle cells (Figures 2E and 2F). At 18.5, Dusp10 expression was noted in the enamel organ, in the ectomesenchymal cells of the dental papillae and the dental follicle cells (Figures 2G and 2H). Later, at P0, Dusp10 expression was localized to the preameloblasts, odontoblasts and pre-dentin (Figures 2I and 2K). Marked expression was noted in the nuclei and cytoplasms of preameloblasts, as well as in the subjacent odontoblast layer (Figure 2K). *Dusp10* mRNA expression was observed in the mouse craniofacial region and oral cavity at E14.5, particularly in the mouth and tongue.(Hoffman et al. 2008) (Figure 2L). These data suggested that Atf1 and Dusp10 are present throughout tooth development stages in mouse embryos and thus likely to have a role in tooth development. Therefore, variations that impact the function of ATF1 and DUSP10 in humans are likely to play a contributory role in tooth agenesis.

**Fig. 1.**
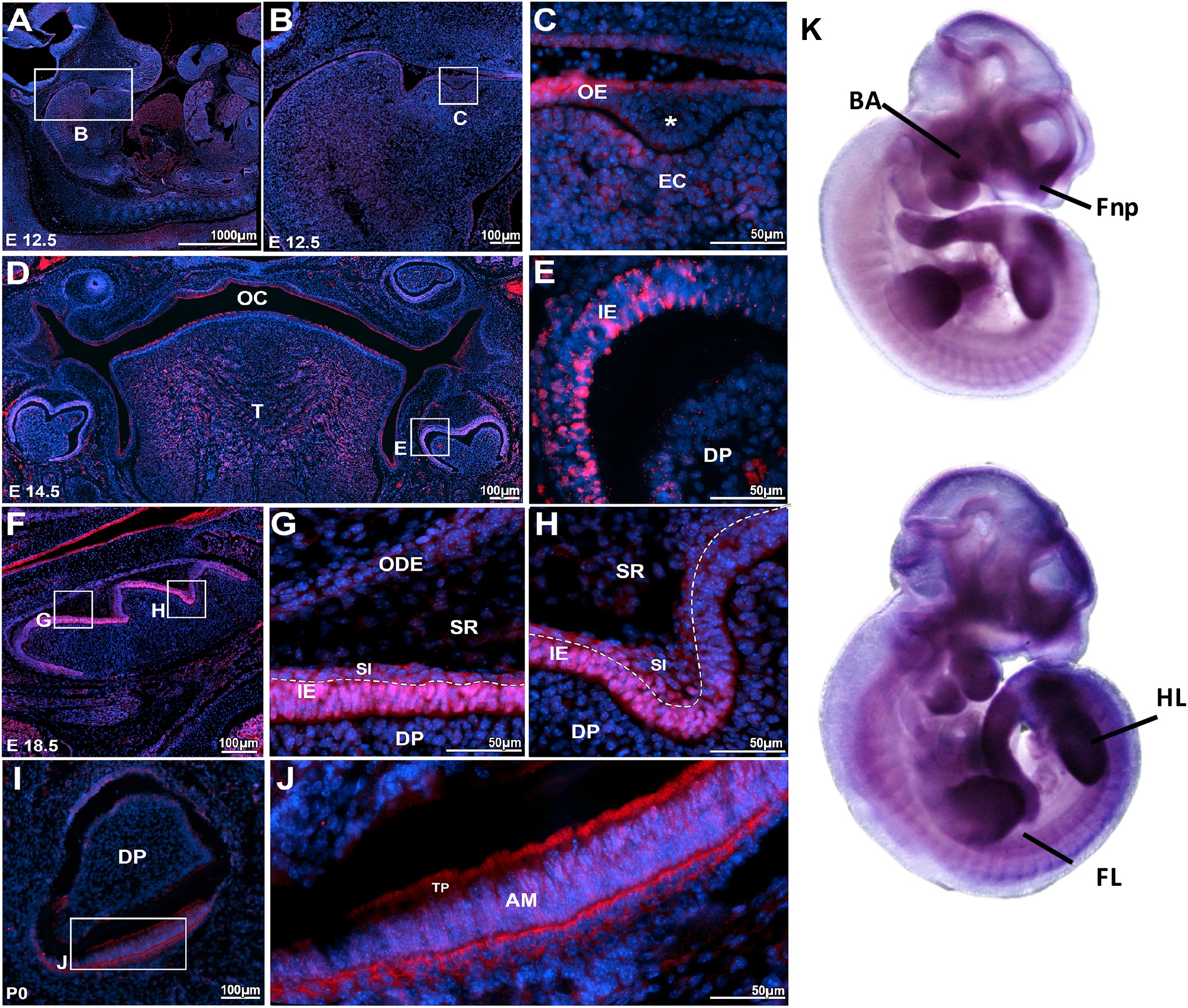
Atf1 is expressed in mouse developing teeth (A, B) Sagittal section of mouse embryo at E12.5. **(C)** Proliferation (*) of the oral epithelium (OE) into the ectomesenchyma, (EM) consistent with the formation of a molar tooth bud at early bud stage. ATF1 expression was detected the oral epithelium, as well as in the condensed ectomesenchymal cells. **(D)** Panoramic photomicrography showing the mouse oral cavity (OC) with tooth germs at early bell stage (E14.5)**. (E)** ATF1 expression was noted in the inner enamel/dental epithelium (IE) and dental papilla (DP). **(F-H)** At E16.5 (Appendix Fig. 1) and E18.5, marked ATF1 expression was observed in the inner enamel epithelium (IE) and in the stratum intermedium (SI), with sparse expression noted in the stellate reticulum (SR) and outer dental (ODE) epithelium. **(I-J)** In incisor teeth at P0, ATF1 expression was particularly evident in the polarized layer of ameloblasts, and in the Tomes’ processes (TP). **(K)** *Atf1* mRNA detected by whole mount in situ hybridization with digoxigenin-labeled antisense RNA followed by alkaline phosphatase-coupled antibody against digoxigenin in C57BL mice at embryonic day 10.5. Strong expression is noted in the mesenchyme of frontonasal prominences, branchial arches and limbs (Obtained from MGI Gene Expression Database. Original source: Gray et al. Mouse Brain Organization Revealed Through Direct Genome-Scale TF Expression Analysis. Science. 2004 24;306(5705):2255-2257). Secondary antibody goat anti-rabbit-Alexa 555 for detection of ATF1 and DAPI for nuclear staining. OE = Oral ephitelium; EM = ectomesenchymal cells; OC = oral cavity; T = tongue; IE = inner epithelium; DP = dental papilla; SI= stratum intermedium; SR= stellate reticulum; ODE outer dental ephithelium; AM = ameloblasts; TP = Tomes’ processes; BA = branchial arches; Fnp = frontonasal processes; HL = hind limbs; FL = forward limbs.

**Fig. 2.**
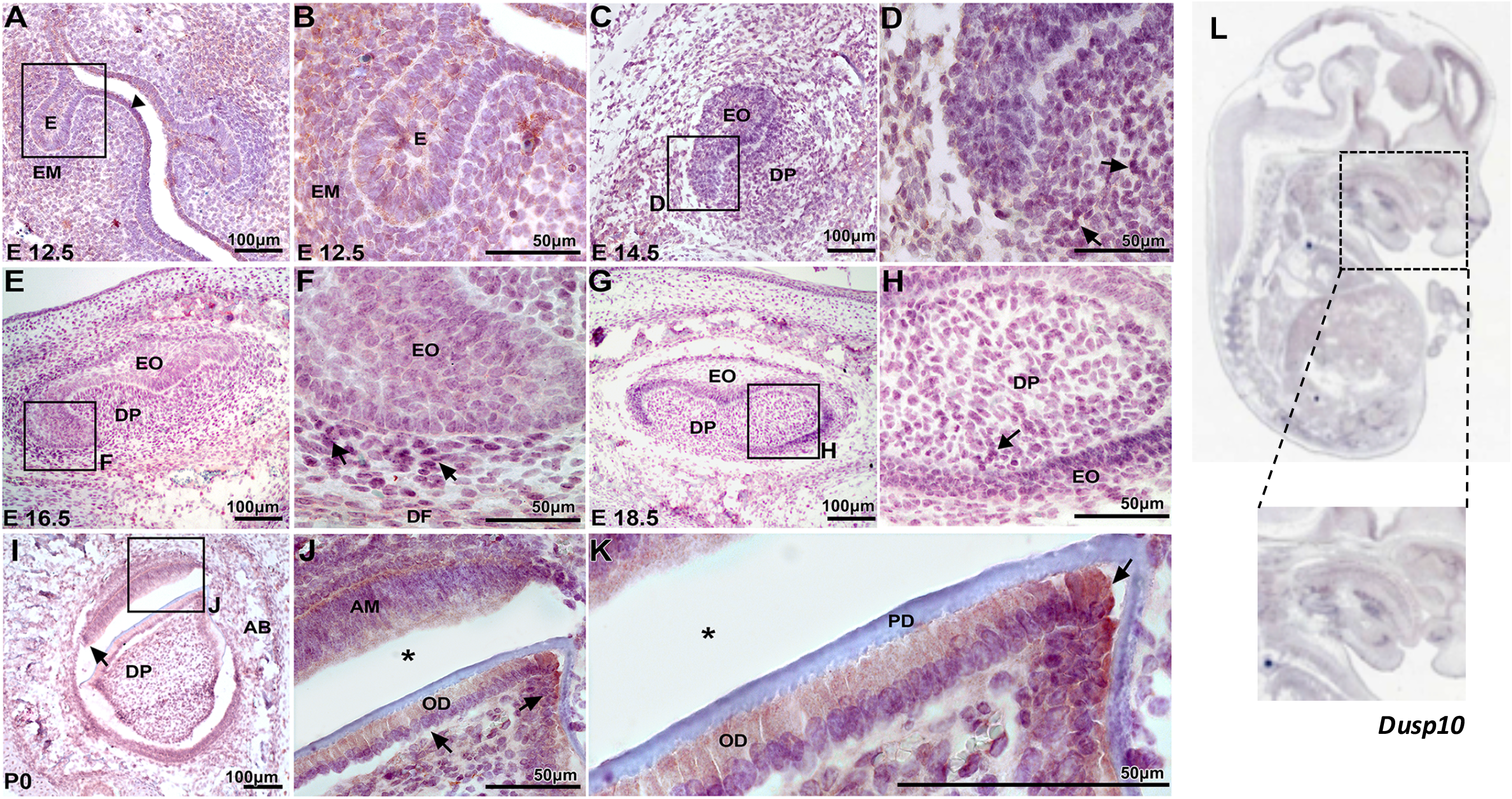
Dusp10 expression in mouse developing teeth (A-B) At E12.5, there is a proliferation of the dental lamina surrounded by a condensation of the underlying ectomesenchymal cells (EM) compatible with a bud stage of tooth development. Dusp10 expression was noted in the epithelial cells from oral epithelium (arrowhead), as well as in the condensed ectomesenchymal cells surrounding the epithelial proliferation (E). **(C,D)** At E14.5, expression was noted detected in the enamel organ (EO) and dental papillae (DP) (arrows). **(E,F)** At bell stage E16.5, Dusp10 positive staining is observed in the enamel organ, dental papillae and dental follicle (DF) cells (arrows). **(G, H)** At E18.5, the tooth germ demonstrates morphodifferentiation compatible with a bell stage. The enamel organ (EO) cells, ectomesenchymal cells of the dental papillae (DP) and the dental follicle cells show homogenous nuclear immune staining. **(I-K)** At P0, the incisor tooth germ is surrounded by alveolar bone (AB) and shows morphology compatible with a late bell stage or crown stage, such as ameloblasts (AM), odontoblasts (OD) and pre-dentin (PD) deposition. Dusp10 expression was observed in the preameloblasts and subjacent odontoblast layer (arrows). **(L)** *Dusp10* mRNA detected by whole mount in situ hybridization in C57BL/6 mouse at E14.5. Positive expression is noted in the craniofacial region and oral cavity, particularly in the mouth and tongue. (Obtained from MGI Gene Expression Database. Original source: Hoffman et al. Genome Biol 2008;9(6):R99). An artifact (*) separated the preameloblasts from the pre-dentin and odontoblastic layer. DAB chromogen and counterstaining with Mayer’s Hematoxylin. OE = Oral ephitelium; EM = ectomesenchymal cells; EO = enamel organ; DP = dental papilla; AM = ameloblasts; AB = alveolar bone.

*CASC8* has only recently been annotated in the human genome, and not yet in the mouse genome, and thus limited our analysis to evaluate its expression in relevant tissues. Using information available in public databases, we found that *CASC8* mRNA is expressed in many tissues including brain and skin, and over-expression was noted in minor salivary gland, colon and esophagus. We then performed RT-PCR analysis of *CASC8* expression in stem cells of the apical papilla (SCAP), a dental-derived stem cell line, (Liu et al. 2006) however, no expression was detected.

## Discussion

Over a decade ago, Lammi et al. (Lammi et al. 2004) identified *AXIN2* as a novel gene involved in tooth development after finding mutations in this gene to be co-segregating with CRC and TA in a large multiplex family. Moreover, these authors showed the expression of *Axin2* in developing mouse teeth and showed for the first time that a gene involved in CRC could also be involved in tooth development. Here, we report the unbiased discovery of new CRC-predisposing genes with a role in tooth development as well as additional candidate variants for isolated TA.

Our association analyses revealed genome-wide significant associations between TA and loci on 1q41, 8q24, and 12q13, containing *DUSP10* (dual specificity phosphatase 10), *CASC8* (cancer susceptibility candidate 8), and *ATF1* (activating transcription factor 1) genes, respectively.

*ATF1* belongs to the ATF/CREB family of transcription factors, which binds to the consensus ATF/CRE site ‘TGACGTCA’ and regulates the transcription of target genes to participate in various cellular processes.(Hai and Hartman 2001) Previous reports showed that the function of *ATF1* appears to depend on the cellular and genetic context to play an important role in tumor progression in a tumor-specific manner.(Huang et al. 2012) ATF1 was found to be over-expressed in several cancer types including lymphoma, melanoma and nasopharyngeal carcinoma, and functioned as a tumor promoter both *in vitro* and *in vivo.*(Hsueh and Lai 1995; Huang et al. 2016; Jean and Bar-Eli 2000; Su et al. 2011) In contrast, disruption of ATF1 activity suppressed its tumorigenicity and metastatic potential.(Jean and Bar-Eli 2000) However, the exact role of *ATF1* during craniofacial development is still largely unknown. Loss of *Atf1* in mice did not cause any obvious phenotypic abnormalities although led to increased programmed cell death.(Bleckmann et al. 2002)

DUSP10 is a dual specificity phosphatase (DUSP) that negatively regulates the activation of the mitogen-activated protein kinase (MAPK) gene family, which includes the extracellular-regulated kinases (ERKs) and the stress-activated protein kinases p38 and c-Jun NH_2_-terminal kinase (JNK). Activation of MAPK by various stimuli including growth factors, cytokines, or stress conditions, regulates major cellular responses such as proliferation, differentiation, survival, migration or production of soluble factors.(Bermudez et al. 2010) *Dusp10* has been regarded as an important regulator of tumorigenesis in animal models,(Png et al. 2016) and loss of *Dusp10* in mice caused enhanced immune and inflammatory responses although effects on skeletal phenotype have not been described.(Zhang et al. 2004) Nonetheless, other DUSPs have been found to play critical roles in development. Loss of function studies revealed that Dusp4 is essential for early development and endoderm specification in the zebrafish,(Brown et al. 2008) whereas Dusp5 was described to control angioblast populations in the lateral plate mesoderm.(Pramanik et al. 2009)

*CASC8* is a long noncoding RNA (lncRNA), located in the gene desert region on 8q24.21. LncRNA is a new class of transcripts that are involved in multiple cellular functions including the regulation of expression of multiple genes. Studies have shown that SNPs in lncRNAs may affect the biological processes of messenger RNA conformation, and results in the modification of its interacting partners. Interestingly, various cancer types, including CRC, prostate, breast and gastric cancer, have been reported in association with *CASC8,* (Lemire et al. 2015; Yao et al. 2015) possibly through its function as a lncRNA.

Variants in *ATF1, DUSP10 and CASC8* have been associated with increased risk of CRC in different GWAS with multiple populations, (Houlston et al. 2010; Hsu et al. 2015; Lemire et al. 2015) although functional studies addressing the potential effects of these variants on gene function and/or disease mechanisms are scarce. While the biological function of the associated *ATF1, DUSP10,* and *CASC8* variants is yet to be determined, our bioinformatics analyses revealed that all three variants are predicted to be within DNAse I hypersensitivity clusters, or in enhancer histone marks, with likely functional effects on gene expression. Moreover, *ATF1* rs11169552 is predicted to harbor a binding site for *POLR2A,* essential for RNA polymerase activity and DNA transcription. Another regulatory variant in *ATF1* (rs11169571) was shown to increase breast/ovarian cancer risk through modifying miRNA binding.(Kontorovich et al. 2010; Yang et al. 2015) Interestingly, this variant is in linkage disequilibrium with *ATF1* rs11169552, associated in the present study, and could be transmitting the same genetic information as well as functional potential. *DUSP10* rs6687758 is predicted to have a putative binding motif for TBP (TATA-box binding protein), which controls the transcription machinery. *CASC8* rs10505477 has a putative binding motif to NK2-5, a homeobox-containing transcription factor with roles in embryonic development. These findings suggest that these genes are active in early cell processes being important in both embryogenesis and tumorigenesis.

The relevance of *ATF1, DUSP10* and *CASC8* in tooth development is unknown. Therefore, we assessed the expression of these genes or their encoded proteins in relevant tissues or cell lines. During mouse embryonic development, *Atf1* mRNA was detected at embryonic day 10.5, with strong expression noted in the mesenchyme of the frontonasal prominences, branchial arches and limbs (Gray et al. 2004). Our findings showed that Atf1 is expressed throughout mouse tooth development stages, in the oral epithelial cells and subjacent condensed ectomesenchymal cells during the early bud stage of tooth development, then shifting to the inner dental epithelium and dental papilla, and finally in the ameloblasts at the final stages of tooth development. *Dusp10* mRNA expression was also observed in the mouse craniofacial region and oral cavity at E14.5, particularly in the mouth and tongue. (Hoffman et al. 2008) In our study, the expression of Dusp10, albeit weak, was also noted in the proliferating dental tissues at the early stages of tooth development. At later stages, Dusp10 expression was evident in the enamel organ, pre-ameloblasts and odontoblastic layer. Expression of *CASC8* mRNA has been observed in the brain and skin, and over-expression noted in minor salivary gland, colon and esophagus. However, no expression of *CASC8* was detected in human dental stem cells of the apical papilla (SCAP). Since *CASC8* has only recently been annotated in the human genome, we were limited in the choice of relevant tissues to assess its expression thus hampering our interpretation of the role of this gene in tooth development. Of note, the observed association between TA and the *CASC8* locus on chromosome 8q24.21 raises intriguing questions as this locus has been associated with increased risk of several malignancies including CRC (Lemire et al. 2015; Yao et al. 2015) and also cleft lip/palate,(Birnbaum et al. 2009; Yu et al. 2017) for which an expanded phenotype including tooth agenesis has been proposed.(Letra et al. 2007)

To date, no large-scale, unbiased genetic studies have been conducted to elucidate the full spectrum of common variants associated with isolated TA, some of which may be located in genes/pathways yet unknown for a role in tooth development. Our findings provide evidence that unbiased approaches have the potential to elucidate the full spectrum of variants associated with TA. It is possible that the associated variants in the present study may be in linkage disequilibrium with a distant causal variant, and acting as surrogate markers for the condition. Additional fine-mapping around the associated loci may provide additional insights into the association signal. Finally, while the exact contributions of *ATF1, DUSP10* and *CASC8* in tooth agenesis remain to be further elucidated, our findings further support that genes involved in colorectal cancer may also be involved in tooth development and provide additional insights into deciphering the complex etiology of the condition.

## Acknowledgements

We thank the study individuals for participation in this study. This work was supported in part by funding from the National Institute for Dental and Craniofacial Research (NIDCR) (R03-DE024596 to A.L., and R90 DE024296-03 to M.R.B.); American Association of Orthodontists (Foundation Biomedical Research Award to S.A. and A.L.); the Rolanette and Berdon Lawrence Bone Disease Program of Texas (to W.D.F.), and the Scientific and Technological Research Institution of Turkey, TUBITAK-ERA NET (CRANIRARE-2, grant number: SBAG-112S398), Istanbul University Research Fund (Project No: 48398). M.W.W. was supported by the UTSD Summer Research Program. Data and samples from Pittsburgh were sourced through the Dental Registry and DNA Repository project, which is supported by the University of Pittsburgh School of Dental Medicine.

## Supplementary Material

Supplementary Material is available online.

## References

Bermudez O, Pages G, Gimond C (2010) The dual-specificity MAP kinase phosphatases: critical roles in development and cancer Am J Physiol Cell Physiol 299:C189–202 doi:10.1152/ajpcell.00347.2009

Birnbaum S et al. (2009) Key susceptibility locus for nonsyndromic cleft lip with or without cleft palate on chromosome 8q24 Nat Genet 41:473–477 doi:10.1038/ng.333

Bleckmann SC, Blendy JA, Rudolph D, Monaghan AP, Schmid W, Schutz G (2002) Activating transcription factor 1 and CREB are important for cell survival during early mouse development Mol Cell Biol 22:1919–1925

Brown JL, Snir M, Noushmehr H, Kirby M, Hong SK, Elkahloun AG, Feldman B (2008) Transcriptional profiling of endogenous germ layer precursor cells identifies dusp4 as an essential gene in zebrafish endoderm specification Proc Natl Acad Sci U S A 105:12337–12342 doi:10.1073/pnas.0805589105

Callahan N, Modesto A, Meira R, Seymen F, Patir A, Vieira AR (2009) Axis inhibition protein 2 (AXIN2) polymorphisms and tooth agenesis Arch Oral Biol 54:45–49 doi:10.1016/j.archoralbio.2008.08.002

Cudney SM, Vieira AR (2012) Molecular factors resulting in tooth agenesis and contemporary approaches for regeneration: a review Eur Arch Paediatr Dent 13:297–304

Gray PA et al. (2004) Mouse brain organization revealed through direct genome-scale TF expression analysis Science 306:2255–2257 doi:10.1126/science.1104935

Hai T, Hartman MG (2001) The molecular biology and nomenclature of the activating transcription factor/cAMP responsive element binding family of transcription factors: activating transcription factor proteins and homeostasis Gene 273:1–11

Hennekam RCK, I.D.; Allanson, J.E. (2010) Gorlin’s Syndromes of the Head and Neck. 5th edn. Oxford University Press, New York, NY

Hoffman BG et al. (2008) Identification of transcripts with enriched expression in the developing and adult pancreas Genome Biol 9:R99 doi:10.1186/gb-2008-9-6-r99

Houlston RS et al. (2010) Meta-analysis of three genome-wide association studies identifies susceptibility loci for colorectal cancer at 1q41, 3q26.2, 12q13.13 and 20q13.33 Nat Genet 42:973–977 doi:10.1038/ng.670

Hsu L et al. (2015) A model to determine colorectal cancer risk using common genetic susceptibility loci Gastroenterology 148:1330–1339 e1314 doi:10.1053/j.gastro.2015.02.010

Hsueh YP, Lai MZ (1995) Overexpression of activation transcriptional factor 1 in lymphomas and in activated lymphocytes J Immunol 154:5675–5683

Huang GL et al. (2012) Activating transcription factor 1 is a prognostic marker of colorectal cancer Asian Pac J Cancer Prev 13:1053–1057

Huang GL et al. (2016) The protein level and transcription activity of activating transcription factor 1 is regulated by prolyl isomerase Pin1 in nasopharyngeal carcinoma progression Cell Death Dis 7:e2571 doi:10.1038/cddis.2016.349

Jean D, Bar-Eli M (2000) Regulation of tumor growth and metastasis of human melanoma by the CREB transcription factor family Mol Cell Biochem 212:19–28

Kang BW et al. (2015) Association between GWAS-identified genetic variations and disease prognosis for patients with colorectal cancer PLoS One 10:e0119649 doi:10.1371/journal.pone.0119649

Kontorovich T, Levy A, Korostishevsky M, Nir U, Friedman E (2010) Single nucleotide polymorphisms in miRNA binding sites and miRNA genes as breast/ovarian cancer risk modifiers in Jewish high-risk women Int J Cancer 127:589–597 doi:10.1002/ijc.25065

Lammi L et al. (2004) Mutations in AXIN2 cause familial tooth agenesis and predispose to colorectal cancer Am J Hum Genet 74:1043–1050 doi:10.1086/386293

Lemire M et al. (2015) A genome-wide association study for colorectal cancer identifies a risk locus in 14q23.1 Hum Genet 134:1249–1262 doi:10.1007/s00439-015-1598-6

Letra A, Menezes R, Granjeiro JM, Vieira AR (2007) Defining subphenotypes for oral clefts based on dental development J Dent Res 86:986–991 doi:10.1177/154405910708601013

Letra A, Menezes R, Granjeiro JM, Vieira AR (2009) AXIN2 and CDH1 polymorphisms, tooth agenesis, and oral clefts Birth Defects Res A Clin Mol Teratol 85:169–173 doi:10.1002/bdra.20489

Li X et al. (2014) A model for the molecular underpinnings of tooth defects in Axenfeld-Rieger syndrome Hum Mol Genet 23:194–208 doi:10.1093/hmg/ddt411

Liu H, Gronthos S, Shi S (2006) Dental pulp stem cells Methods Enzymol 419:99–113 doi:10.1016/S0076-6879(06)19005-9

Liu X, Wu C, Li C, Boerwinkle E (2016) dbNSFP v3.0: A One-Stop Database of Functional Predictions and Annotations for Human Nonsynonymous and Splice-Site SNVs Hum Mutat 37:235–241 doi:10.1002/humu.22932

Navarro M, Nicolas A, Ferrandez A, Lanas A (2017) Colorectal cancer population screening programs worldwide in 2016: An update World J Gastroenterol 23:3632–3642 doi:10.3748/wjg.v23.i20.3632

Png CW et al. (2016) DUSP10 regulates intestinal epithelial cell growth and colorectal tumorigenesis Oncogene 35:206–217 doi:10.1038/onc.2015.74

Polder BJ, Van’t Hof MA, Van der Linden FP, Kuijpers-Jagtman AM (2004) A meta-analysis of the prevalence of dental agenesis of permanent teeth Community Dent Oral Epidemiol 32:217–226 doi:10.1111/j.1600-0528.2004.00158.x

Pramanik K et al. (2009) Dusp-5 and Snrk-1 coordinately function during vascular development and disease Blood 113:1184–1191 doi:10.1182/blood-2008-06-162180

Purcell S et al. (2007) PLINK: a tool set for whole-genome association and population-based linkage analyses Am J Hum Genet 81:559–575 doi:10.1086/519795

Ranade K et al. (2001) High-throughput genotyping with single nucleotide polymorphisms Genome Res 11:1262–1268 doi:10.1101/gr.157801

Siegel R, Desantis C, Jemal A (2014) Colorectal cancer statistics, 2014 CA Cancer J Clin 64:104–117 doi:10.3322/caac.21220

Su B et al. (2011) Stage-associated dynamic activity profile of transcription factors in nasopharyngeal carcinoma progression based on protein/DNA array analysis OMICS 15:49–60 doi:10.1089/omi.2010.0055

Thesleff I, Sharpe P (1997) Signalling networks regulating dental development Mech Dev 67:111–123

Thesleff I, Vaahtokari A, Vainio S, Jowett A (1996) Molecular mechanisms of cell and tissue interactions during early tooth development Anat Rec 245:151–161 doi:10.1002/(SICI)1097-0185(199606)245:2<151::AID-AR4>3.0.CO;2-#

Yang S et al. (2015) Association of single nucleotide polymorphisms in the 3’UTR of ERAP1 gene with essential hypertension in the Northeastern Han Chinese Gene 560:211–216 doi:10.1016/j.gene.2015.02.005

Yao K, Hua L, Wei L, Meng J, Hu J (2015) Correlation Between CASC8, SMAD7 Polymorphisms and the Susceptibility to Colorectal Cancer: An Updated Meta-Analysis Based on GWAS Results Medicine (Baltimore) 94:e1884 doi:10.1097/MD.0000000000001884

Yin W, Bian Z (2015) The Gene Network Underlying Hypodontia J Dent Res 94:878–885 doi:10.1177/0022034515583999

Yin W, Bian Z (2016) Hypodontia, a prospective predictive marker for tumor? Oral Dis 22:265–273 doi:10.1111/odi.12400

Yu Y et al. (2017) Genome-wide analyses of non-syndromic cleft lip with palate identify 14 novel loci and genetic heterogeneity Nat Commun 8:14364 doi:10.1038/ncomms14364

Zhang B et al. (2014) Large-scale genetic study in East Asians identifies six new loci associated with colorectal cancer risk Nat Genet 46:533–542 doi:10.1038/ng.2985

Zhang Y et al. (2004) Regulation of innate and adaptive immune responses by MAP kinase phosphatase 5 Nature 430:793–797 doi:10.1038/nature02764

